# FOX transcription factors are common regulators of Wnt/β-catenin signaling

**DOI:** 10.1101/2022.12.13.520306

**Authors:** Lavanya Moparthi, Stefan Koch

## Abstract

The Wnt/β-catenin signaling pathway is a critical regulator of development and stem cell maintenance. Mounting evidence suggests that the context-specific outcome of Wnt signaling is determined by the collaborative action of multiple transcription factors, including members of the highly conserved forkhead box (FOX) protein family. However, the contribution of FOX transcription factors to Wnt signaling has not been investigated in a systematic manner. Here, we performed uniform gain-of-function screens of all 44 human FOX transcription factors to identify and classify new regulators of the Wnt/β-catenin pathway. By combining β-catenin reporter assays with Wnt pathway-focused qPCR arrays and proximity proteomics of selected FOX family members, we determine that most FOX proteins are involved in the regulation of Wnt pathway activity and the expression of Wnt ligands and target genes. Moreover, as a proof of principle we characterize class D and I FOX transcription factors as physiologically relevant positive and negative regulators of Wnt/β-catenin signaling, respectively. We conclude that FOX proteins are common regulators of the Wnt/β-catenin pathway that may control the outcome of Wnt signaling in a tissue-specific manner.

## INTRODUCTION

The Wnt/β-catenin signaling pathway, also referred to as the canonical Wnt pathway, is a central homeostatic signaling cascade, whose defining feature is the Wnt ligand-induced stabilization of β-catenin resulting in the differential expression of Transcription factor 7 / Lymphoid enhancer-binding factor (TCF/LEF) target genes (*1*). Wnt signaling is critical for stem cell maintenance and proliferation, and dysregulation of Wnt activity contributes to major diseases such as cancers. The Wnt transcriptional response is mediated predominantly by TCF/LEF transcription factors (*2, 3*). However, mounting evidence suggests that the context-specific outcome of Wnt signaling is determined by the collaborative action of various other transcription factors that control the expression of Wnt target genes (*4–6*).

Among these additional Wnt regulators are forkhead box (FOX) transcription factors. FOX proteins comprise one of the largest transcription factor families, with 44 main family members in humans that are subdivided into 19 classes (A through S) based on sequence similarity (*7–9*). Several lines of evidence across species suggests that there is substantial functional redundancy between FOX proteins that may not be limited to just closely related family members (*10–13*). In the context of Wnt signaling, numerous FOX proteins have been shown to regulate pathway activity by altering Wnt ligand expression, affecting the subcellular shuttling of β-catenin, and changing the composition of the Wnt transcriptional complex, among others (*9, 14, 15*). Accordingly, the role of some FOX proteins in, for example, carcinogenesis or lifespan control has been attributed to their activity in the Wnt pathway (*16, 17*). However, owing to the large number and divergent expression pattern of FOX transcription factors, the extent of their redundancy in Wnt pathway regulation remains poorly understood.

We therefore assessed the impact of the entire FOX transcription factor family on Wnt signaling using complementary, uniform gain-of-function screens in model cell lines. We find that most FOX proteins regulate Wnt-dependent transcription and target gene expression, and identify class D and I FOXs as Wnt signaling activators and inhibitors, respectively. Our findings establish FOX transcription factors as common regulators of Wnt/β-catenin signaling, and provide mechanistic clues for the apparent functional redundancy of some FOX proteins in the Wnt pathway.

## RESULTS

### FOX transcription factors regulate Wnt/β-catenin signaling

To identify potential new Wnt regulators within the FOX family, we performed gain-of-function screens using two complementary standard assays for Wnt pathway activity: the β-catenin/TCF transcriptional reporter TOPflash (*18*), and a qPCR array of multiple well-characterized Wnt target genes (Fig. 1A). In both assays, we induced FOX proteins by transient overexpression of a functionally validated, uniform FOX plasmid library (*19*). TOPflash assays were performed in 293T cells with or without exogenous Wnt3a and R-spondin 3 (i.e., Wnt pathway on or off), as well as in HCT116 colorectal cancer cells with constitutively active Wnt signaling. qPCR assays were done in 293T cells pretreated with R-spondin 3 to increase Wnt receptor levels and thereby sensitize cells to subsequent pathway activation.

**Figure 1:**
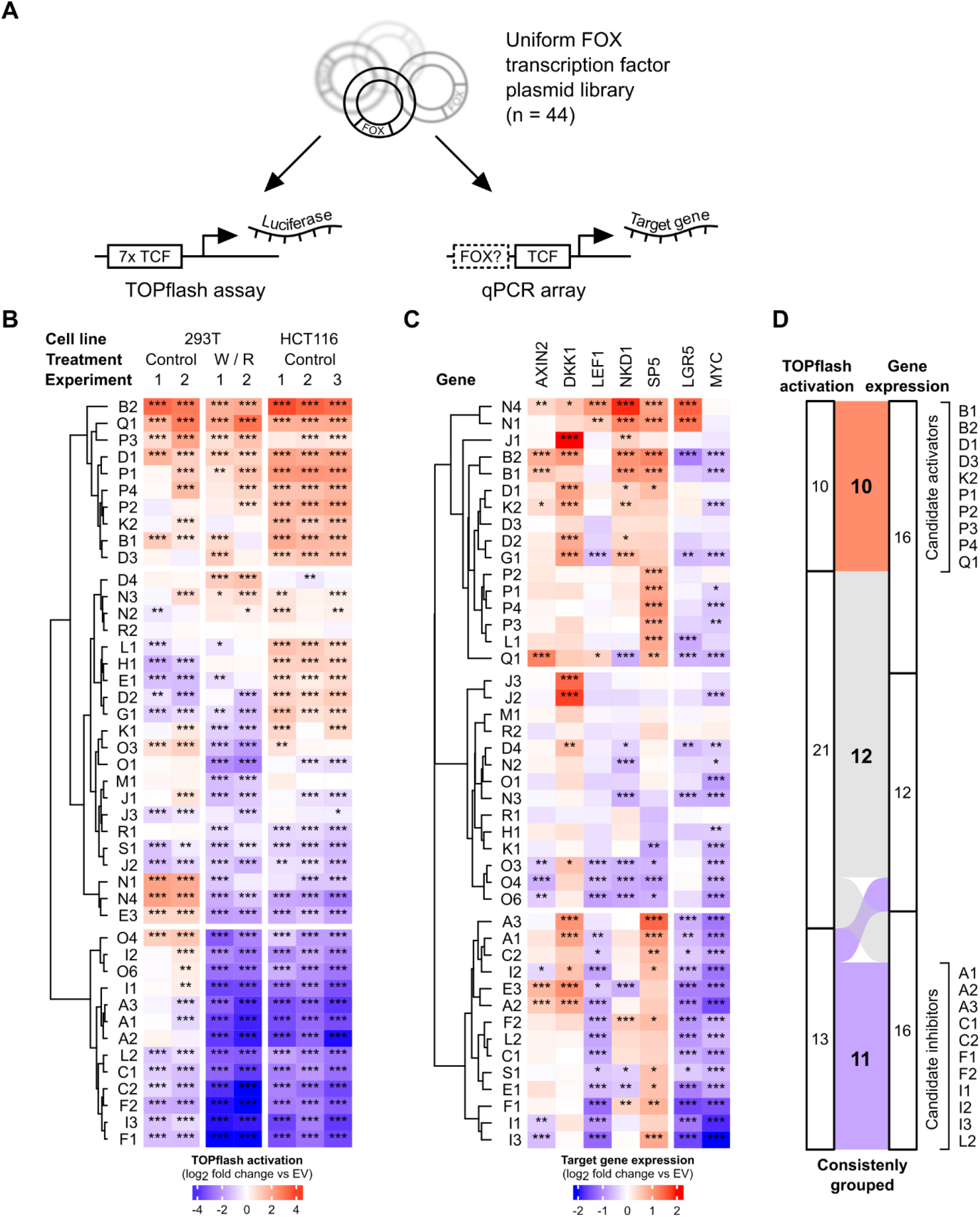
FOX transcription factors are Wnt pathway regulators. (A) Schematic representation of the complementary gain-of-function screens used to identify Wnt pathway regulators in the FOX transcription factor family. (B) Heatmap of β-catenin/TCF reporter (TOPflash) regulation by FOX transcription factors. Where indicated, cells were treated with Wnt3a / R-spondin 3 (W / R) conditioned media. (C) Heatmap of Wnt target gene expression changes following FOX gain-of-function. Gene expression was determined in 293T cells pretreated with 5 ng/ml recombinant human R-spondin 3. (D) Alluvial plot summarizing the grouping of FOX family members based on results from the combined screens. Data in B and C were normalized to the empty vector (EV) control in each column, with each cell showing the average of biological triplicates. Data for each experiment or target gene were analyzed using Dunnett’s post-hoc test against EV following one-way ANOVA (*** *P*<0.001, ** *P*<0.01, * *P*<0.05).

We observed that most FOX transcription factors significantly altered TOPflash activity across multiple independent experiments (Fig. 1B, and supplemental Fig. 1A). We performed unsupervised k-means clustering of the aggregated results to group FOXs into three discrete clusters (Fig. 1B). The first cluster contained several previously identified positive regulators of Wnt signaling, including FOXB2, FOXP1, FOXQ1, and FOXK2 (supplemental Table 1), as well as additional uncharacterized FOX proteins. Similarly, we identified a candidate inhibitor cluster that contained the well-studied negative Wnt pathway regulators FOXO4 and FOXF1/2, alongside others. For validation, we first repeated these experiments using the control plasmid FOPflash (*18*), which was negligibly regulated by most tested FOX proteins (supplemental Fig. 1B). Additionally, we performed TOPflash assays in 293T cells with genetic deletion of LRP5/6, β-catenin, or β-catenin and TCF/LEF (penta-KO) (*20, 21*), which are Wnt refractory (supplemental Fig. 1C, D). In these cells, most FOX proteins had essentially no effect, and did not synergize with Wnt3a / R-spondin 3 in TOPflash activation.

We then performed qPCR arrays of five bona fide β-catenin/TCF target genes in 293T cells (*AXIN2, DKK1, LEF1, NKD1, SP5;* (*21*)), as well as two Wnt target genes that have been studied extensively in other contexts (*LGR5, MYC*) (Fig. 1C, and supplemental Fig. 2A, B). FOX transcription factors can control Wnt target genes independently of β-catenin/TCF (*20, 21*), and we observed considerable variability in the regulation of different Wnt targets. Importantly, however, correlation analysis between the combined gene expression results and TOPflash data (Fig. 1B, C) showed a significant inter-assay correlation (Pearson’s ρ = 0.21; p = 0.0001) that was primarily driven by the differential regulation of *AXIN2, LEF1*, and *MYC* (supplemental Fig. 2C). We thus clustered the gene expression data as before, and observed that most FOX proteins were grouped consistently with the TOPflash data (Fig. 1C, D). Collectively, this approach shortlisted 10 likely activators and 11 candidate inhibitors of Wnt/β-catenin signaling in the FOX family (Fig. 1D), including FOXs that have not been studied in this context previously.

### FOX proteins control the expression of secreted Wnt regulators

We next explored different mechanisms by which FOX transcription factors may regulate Wnt signaling, starting with the induction of Wnt agonists and inhibitors (*15*). We assessed the expression of all 19 Wnt ligands, four R-spondins, and nine secreted Wnt antagonists following FOX induction by qPCR (Fig. 2A, supplemental Fig. 3). We observed that several FOXs induced a group of Wnt agonists that included WNT4, WNT7A/B, and R-spondin 1, among others. Consistently, inhibition of Wnt ligand secretion had a more pronounced effect on TOPflash activation by FOXP1/3/4 compared to FOXP2 and FOXNs, which did not induce Wnts to the same extent (Fig. 2B, C). However, we observed the same pattern of Wnt agonist induction by candidate activators as well as inhibitors, and across all FOX transcription factors, Wnt agonist induction was not generally indicative of TOPflash activation or Wnt target gene expression (Fig. 2D, E). Similarly, while several FOXs differentially regulated various secreted Wnt antagonists (supplemental Fig. 3), we did not observe any pattern consistent with their activity in Wnt pathway assays.

**Figure 2:**
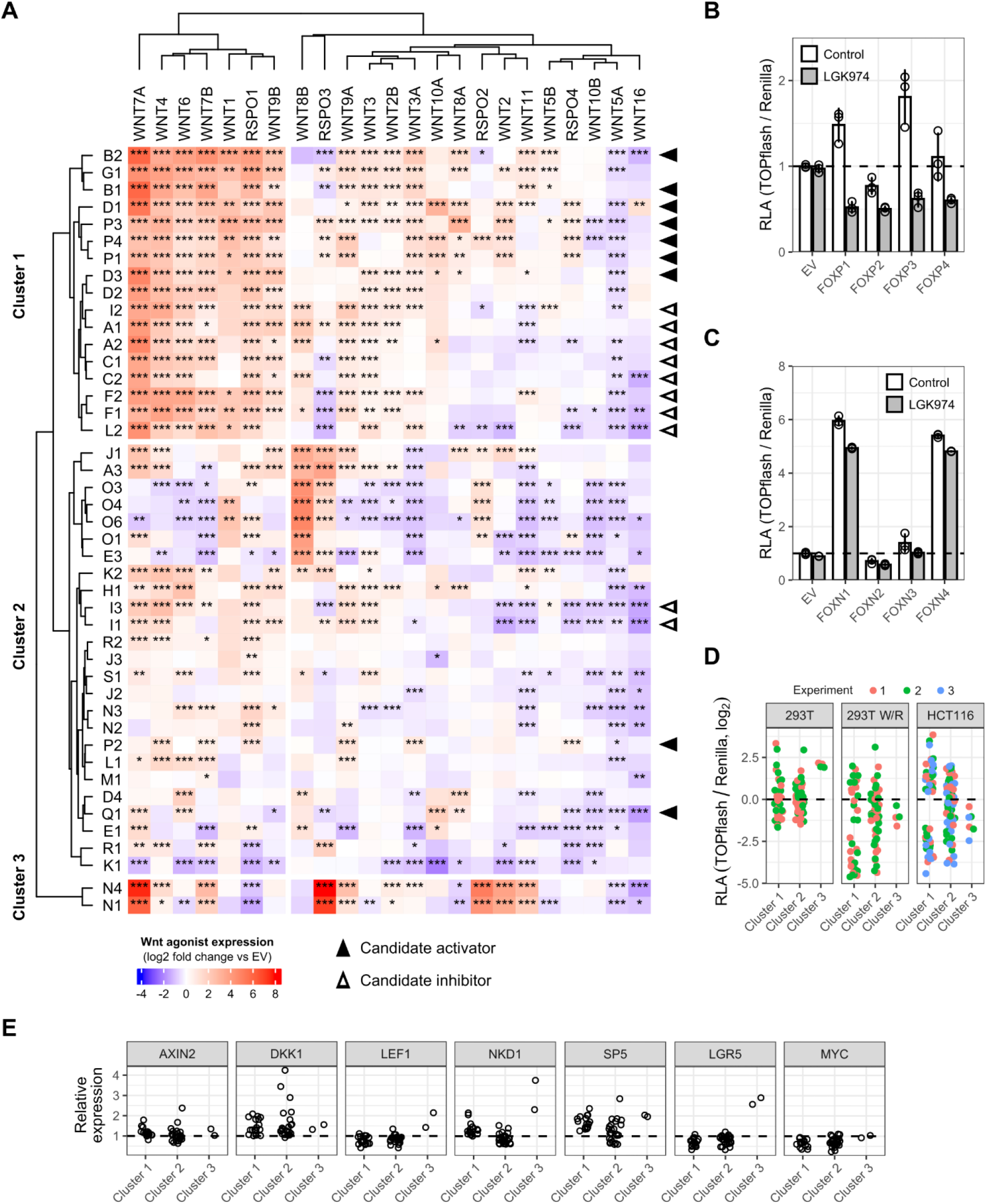
FOX transcription factors regulate Wnt agonist expression. (A) Heatmap of Wnt ligand and R-spondin expression changes following FOX gain-of-function. Gene expression was determined in 293T cells pretreated with 5 ng/ml recombinant human R-spondin 3. Each cell represents the average of three biological replicates, normalized to empty vector (EV) control. (B, C) TOPflash assay in 293T following FOXP (B) or FOXN (C) overexpression. Where indicated, cells were treated with 10 nM LGK974 to block Wnt ligand secretion. (D, E) Grouping of TOPflash (D) and qPCR results (E) from figure 1 by clusters of Wnt agonist expression. Data in panel A target gene were analyzed using Dunnett’s post-hoc test against EV following one-way ANOVA (*** *P*<0.001, ** *P*<0.01, * *P*<0.05). RLA, relative luciferase activity.

We conclude that FOX transcription factors control the expression of Wnt agonists and antagonists, but that this mechanism is insufficient to explain the activity of most family members in the Wnt pathway.

### FOX proteins share a common proximity interactome with TCF/LEF

FOX proteins occupy the promoter regions of various Wnt target genes (*21*), and we have recently shown that FOXQ1 may control gene expression by recruiting similar transcription co-factors as TCF/LEF (*20*). To determine if this is also the case for other FOXs, we used TurboID-based proximity proteomics (*22*) to explore the interaction landscape of five additional FOXs, which displayed widely different transcriptional activity in Wnt and forkhead box reporter assays (*19*). We identified 210 proteins as high-confidence interactors across all six tested FOXs, including our or earlier TurboID-FOXQ1 data (Fig. 3A). Of these, 144 (68%) were associated with four or more FOX proteins, suggesting that they are common FOX interactors. Notably, this number is considerably higher than shared interactors identified in previous co-immunoprecipitation / mass spectrometry assays of 36 FOX transcription factors (*23, 24*) (supplemental Fig. 4), likely due to the better performance of proximity proteomics in screens of chromatin-associated proteins (*25*). Gene ontology analysis of these core interactors revealed a significant enrichment of proteins involved in RNA processing and histone modification, as expected (Fig. 3B). We then asked to what extent the tested FOXs are associated with the same protein complexes as TCF/LEF. For this, we performed an enrichment analysis against a dataset of curated human protein complexes (*26*), including previously identified Tcf7l1, FOXQ1, and FOXB2 interactors (*27, 28*). We observed that FOXQ1, FOXN4, FOXDI, and FOXI1, in particular, share a substantial number of interacting complexes with one another and Tcf7l1 (Fig. 3C, and supplemental Fig. 5). These included chromatin remodeling as well as histone acetylation and methylation complexes known to regulate Wnt target gene expression (*29*). Of particular interest, we found that multiple FOXs were associated with the histone methyltransferase KMT2A, which was recently identified as a critical regulator of Wnt target gene expression (*30*). In contrast, we did not detect association of any FOX family member with β-catenin, TCF/LEF, or BCL9/L.

**Figure 3:**
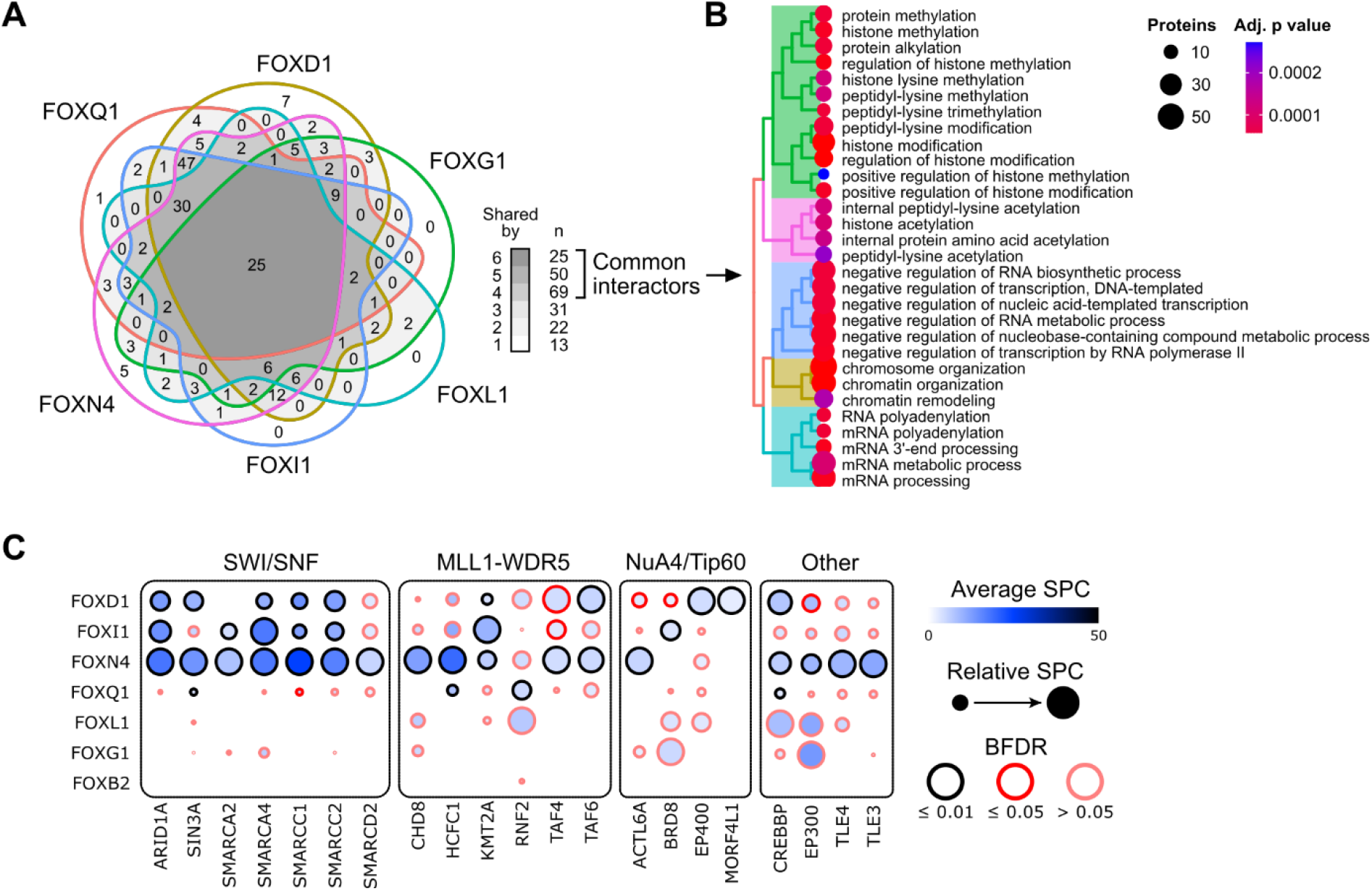
Proximity proteomics identify common and unique FOX interactors. (A) Venn diagram of high-confidence interactors of the indicated FOX proteins identified by TurboID-based proximity proteomics. (B) Gene ontology analysis of biological processes mediated by common FOX interactors. (C) Dot plot of important FOX interactors grouped by functional protein complex, indicated above. Proteomics results were from experiments with two to four biological replicates per bait or corresponding control condition. SPC, spectral count. BFDR, Bayes false discovery rate.

### FOX D and I proteins are novel Wnt pathway regulators

Our results so far suggested widespread control of Wnt signaling by FOX transcription factors. To further validate some of the candidates, we investigated FOXD and FOXI family members, which were recently highlighted as potential regulators of Wnt signaling (*31, 32*). Consistent with our earlier observations, FOXDs activated TOPflash in HCT116 as well as MCF7 breast cancer cells, whereas FOXIs inhibited Wnt reporter activity (Fig. 4A). Moreover, overexpression of FOXD1 and FOXI1 in HCT116 differentially regulated Wnt target gene expression (Fig. 4B). Curiously, contrary to results from 293T, FOXI1 strongly induced *LEF1* in these assays, possibly due to its tissue-specific regulation in colorectal cancer cells (*33*).

**Figure 4:**
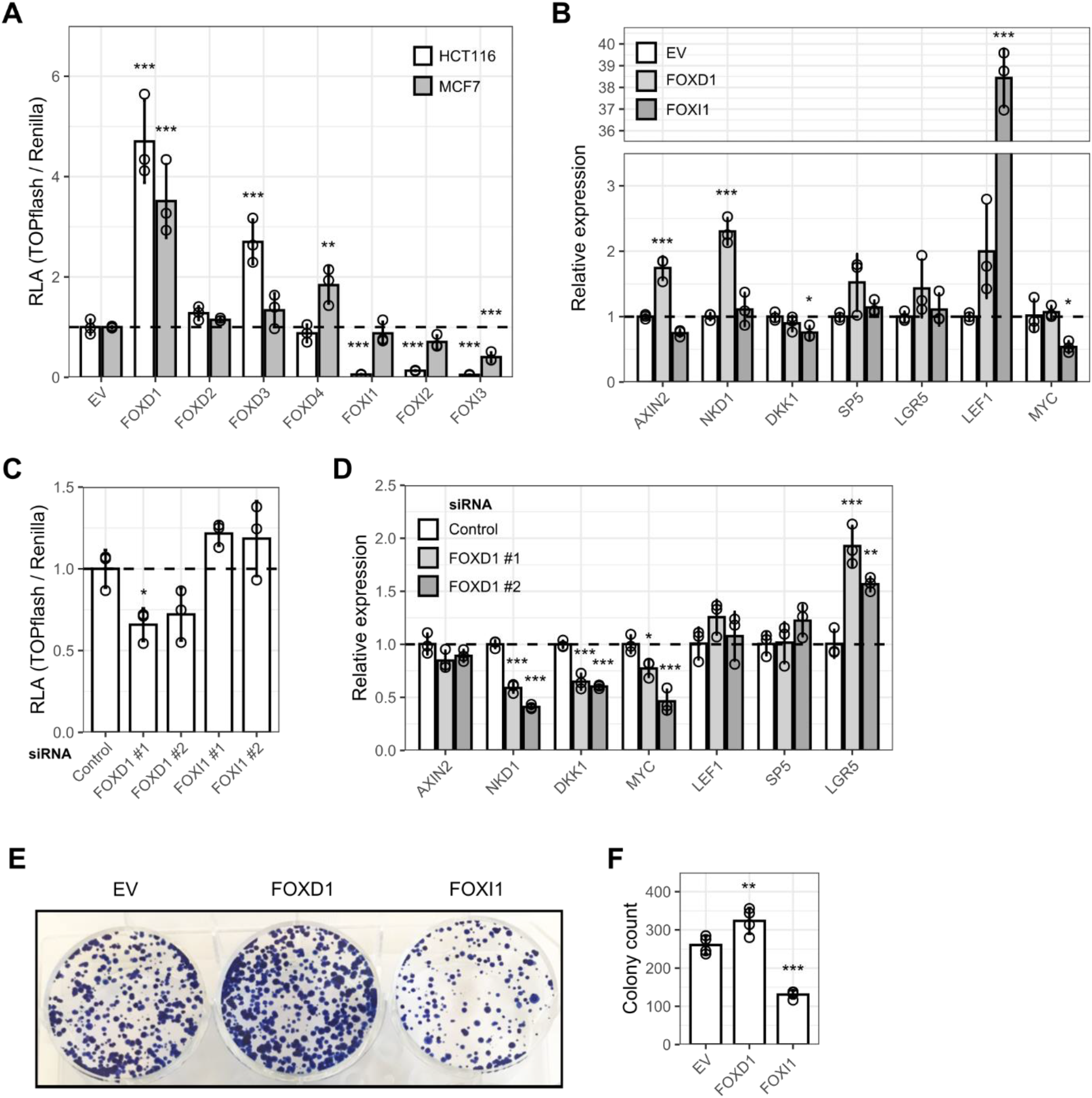
FOXD and FOXI proteins are Wnt pathway regulators. (A) TOPflash assay in HCT116 and MCF7 cells. MCF7 were pretreated with 5 μM CHIR99021. (B) Wnt target gene expression in HCT116 following FOXD1 or FOXI1 expression, as determined by qPCR. (C) TOPflash assay in 293T following FOXD1 or FOXI1 depletion with two independent siRNAs. (D) Wnt target gene expression in 293T following FOXD1 depletion, as determined by qPCR. (E) Representative image of colony formation assays in HCT116 following FOXD1 or FOXI1 gain-of-function. (F) Quantification of colony numbers from n=4 biological replicates. Data in all panels were analyzed using Dunnett’s post-hoc test against EV or siControl following one-way ANOVA (*** *P*<0.001, ** *P*<0.01, * *P*<0.05). RLA, relative luciferase activity.

Conversely, RNA interference of FOXD1 reduced TOPflash activity in 293T, while FOXI1 knock-down increased reporter activity (Fig. 4C). FOXD1 depletion also reduced the expression of multiple Wnt target genes (Fig. 4D), although FOXI1 knock-down had no effect in these assays presumably due to low siRNA efficiency (supplemental Fig. 6). Finally, we determined the effect of FOXD1 and FOXI1 gain-of-function on colony formation in HCT116, which depend on β-catenin/TCF signaling for efficient proliferation (*34*). We observed that FOXD1 and FOXI1 significantly increased and decreased colony formation, respectively (Fig. 4E, F). Taken together, these findings support a physiologically relevant role of class D and I FOXs in the regulation of Wnt signaling.

### FOXD1 and FOXI1 have discrete modes of action in the Wnt pathway

Given that FOXD1-3 strongly induced the expression of multiple Wnt ligands, we asked if this mechanism may sufficiently explain their activity, as we have previously shown for FOXB2 (*28*). Consistent with this hypothesis, TOPflash activation by FOXDs was blocked by LGK974 treatment (Fig. 5A). Moreover, FOXD1-3 were unable to activate TOPflash in 293T ΔLRP5/6 cells expressing constitutively active LRP6 lacking its ligand-binding extracellular domain (Fig. 5B). Interestingly, FOXD4 still activated TOPflash in this assay, suggesting that its mode of action is distinct. Because WNT7A/B were among the most highly induced Wnt ligands, we depleted WNT7B or the WNT7 co-receptor RECK by RNA interference. Under these conditions, FOXD1 was unable to activate TOPflash in 293T (Fig. 5C). Additionally, FOXD1 and FOXD3 in particular were able to modestly activate WNT7B and WNT1 promoter reporters, consistent with direct transcriptional regulation of these ligands (Fig. 5D). Taken together, these results suggest that FOXD1-3 activate Wnt signaling by inducing agonistic Wnt ligands.

**Figure 5:**
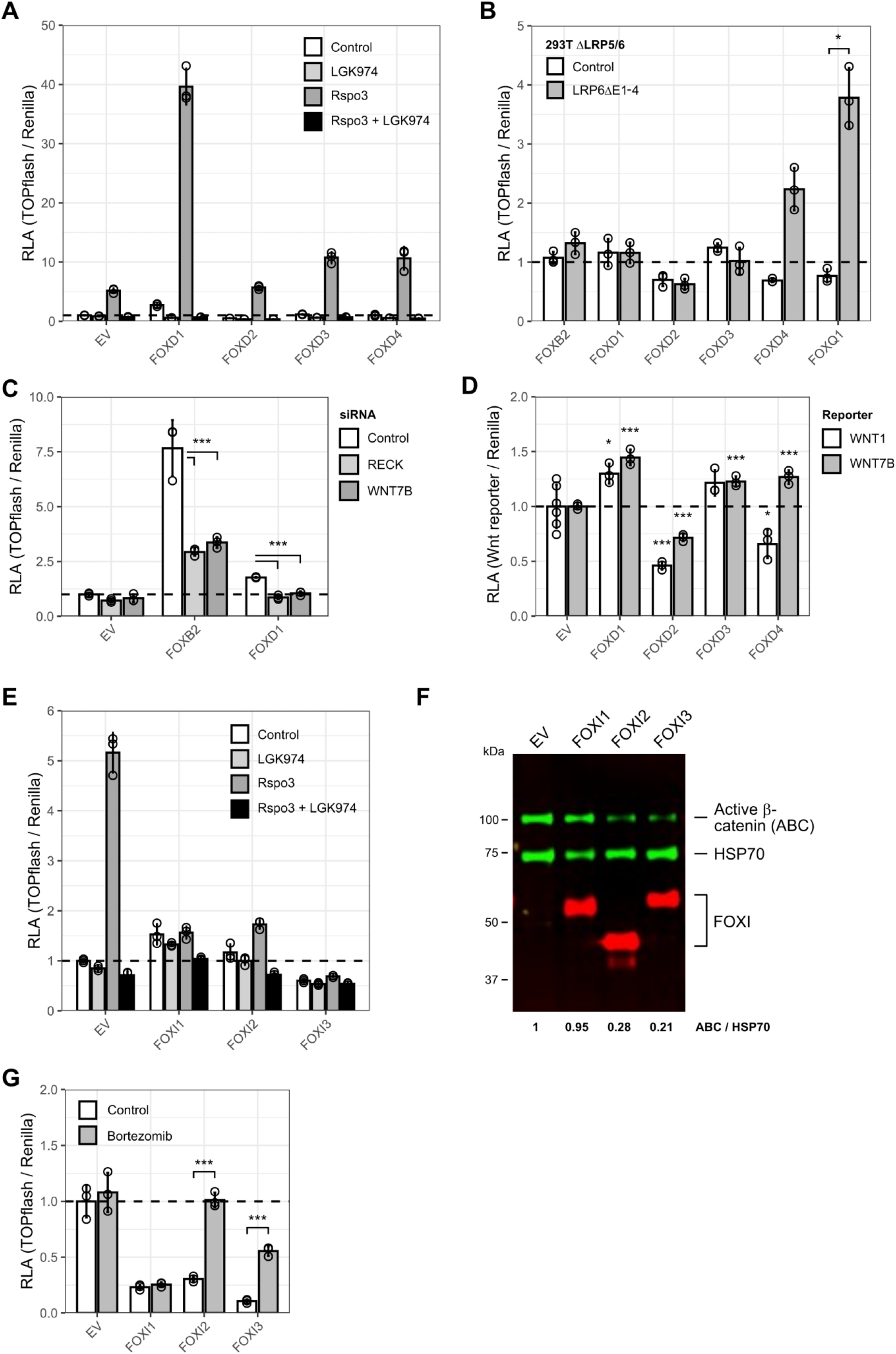
FOXD1 and FOXI1 have discrete functions in the Wnt pathway. (A) TOPflash assay in 293T. Where indicated, cells were treated with 5 ng/ml recombinant human R-spondin 3 (Rspo3) or 10 nM LGK974. (B) TOPflash assay in 293T ΔLRP5/6. Where indicated, cells were transfected with constitutively active LRP6 (LRP6ΔE1-4) to uncouple Wnt activation from Wnt ligand binding. FOXQ1 and FOXB2 were included as positive and negative controls, respectively. Data were normalized to the respective empty vector control. (C) TOPflash assay in 293T following WNT7B or RECK depletion. (D) WNT1 and WNT7B promoter reporter assay in 293T. (E) TOPflash assay in 293T. Where indicated, cells were treated with 5 ng/ml recombinant human R-spondin 3 (Rspo3) or 10 nM LGK974. (F) Immunoblot of active β-catenin (ABC) levels in Wnt3a-treated 293T following FOXI expression. The relative ratio of ABC versus HSP70 housekeeping control is indicated below. (G) TOPflash assay in HCT116. Where indicated, cells were treated with 100 nM proteasome inhibitor bortezomib. Data in B and G were analyzed using an unpaired Welch’s t-test with Bonferroni-Hochberg correction for multiple testing. Data in C and D were analyzed using Dunnett’s post-hoc test against EV or siControl following one-way ANOVA (*** *P*<0.001, * *P*<0.05). RLA, relative luciferase activity.

FOXIs, on the other hand, had a limited impact on Wnt ligand expression, and their inhibitory activity was not affected by LGK974 treatment (Fig. 5E). Several FOX family members such as FOXOs inhibit Wnt signaling by competitive binding of β-catenin (*35*). We observed that FOXIs inhibited TOPflash activation following β-catenin stabilization or overexpression (supplemental Fig. 7A, B). However, we did not detect physical interaction of FOXIs with β-catenin (supplemental Fig. 7C), arguing against β-catenin sequestration. Instead, we noted a destabilization of β-catenin protein by FOXIs. (Fig. 5F). Consistently, proteasomal inhibition released the effect of at least FOXI2/3 on TOPflash activity in HCT116 (Fig. 5G). We conclude that FOXIs may inhibit Wnt signaling in part by promoting the degradation of β-catenin.

FOXD1 and FOXI1 have opposing functions in the Wnt pathway despite a largely overlapping interactome and virtually identical DNA recognition motifs (supplemental Fig. 8A). It is thus likely that their effects are mediated by few specific co-factors. Pairwise analysis of the TurboID data confirmed that most identified interactors were shared between FOXD1 and FOXI1 (Fig. 6A). Accordingly, pathway-focused analyses did not reveal any significant results due to the low number of unique hits. However, among the FOXD1-exclusive hits we detected the ubiquitin/SUMO ligases PIAS4 and TRIM33 (Fig. 6B), which have been implicated in the regulation of Wnt signaling (*36–38*). We also noted that FOXD1 specifically interacted with JUNB (supplemental Fig. 8B), which has been shown to control the expression of multiple Wnt ligands including WNT7A/B, and which was also identified as a candidate FOXB2 interactor (*28, 39*). In contrast, FOXI-exclusive interactors included several transcription co-factors such as BCORL1 and ARID3A (Fig. 6C), which may conceivably mediate its specific effects.

**Figure 6:**
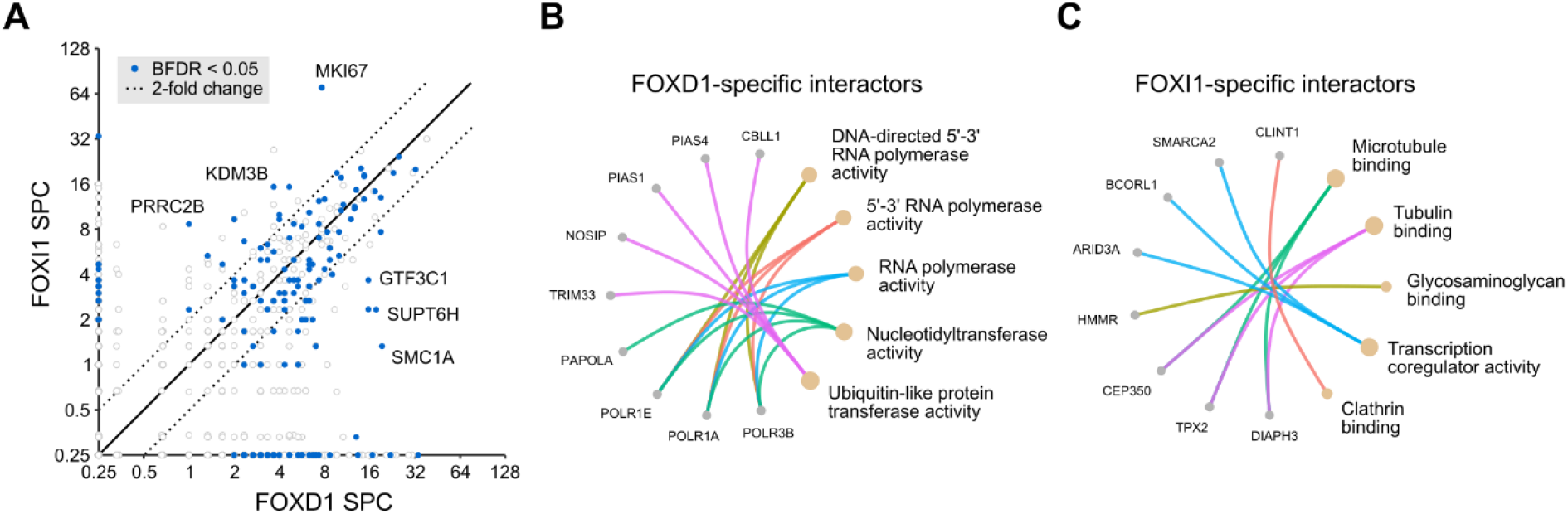
FOXD1 and FOXI1 recruit distinct transcription co-factors. (A) Scatter plot of candidate interactors of FOXD1 and FOXI1 identified by TurboID proximity proteomics. Some interactors that are enriched in one of the samples are highlighted. Hits along the axes were considered specific interactors. (B, C) Functional grouping of specific interactors of (B) FOXD1 and (C) FOXI1, based on gene ontology molecular function terms. SPC, average spectral count; BFDR, Bayes false discovery rate.

## DISCUSSION

Numerous reports have implicated FOX transcription factors in the regulation of Wnt/β-catenin signaling (*15*), but mainly due to inconsistent methodology it is difficult to draw overarching conclusions from these earlier studies. Using complementary gain-of-function assays of all human FOXs, we now show that most FOX proteins affect Wnt transcriptional activity, target gene expression, and Wnt ligand induction, including family members that have not been investigated in this context. Moreover, we show that multiple FOX proteins associate with similar transcription co-factor complexes as Tcf7l1, and may thus act as β-catenin-independent regulators of Wnt target genes. Finally, we use these observations to characterize FOXD and FOXI transcription factors as Wnt pathway regulators.

In recent years, there has been a renewed interest in investigating the transcriptional regulation of Wnt target genes, many of which have important functions in development and pathobiology (*29*). Although it is thought that most of these target genes are controlled primarily by TCF/LEF, other transcription factors such as TBX, CDX, and SOX family members may contribute to the regulation of specific Wnt targets (*4–6*). FOX transcription factors are similarly known to control Wnt pathway activity in various contexts, and it has been shown that several prototypical Wnt targets including *AXIN2* are differentially regulated by FOXs independently of β-catenin/TCF (*20, 21, 28*). Our findings expand on these observations, and highlight new candidate Wnt regulators within the FOX family. For example, we show that FOXO6 affects Wnt reporter activity and target gene expression to a similar extent as other FOXO proteins. Although this finding is not entirely surprising considering that FOXO3/4 in particular are well-established Wnt inhibitors (*17, 35, 40*), the role of FOXO6 remained to be formally established. FOXO6 has distinct subcellular shuttling dynamics and a more restricted expression pattern compared to other FOXOs (*9, 41*), and contributes to the pathogenesis of several types of tumors, including breast and colorectal cancer (*42, 43*). It thus appears worthwhile to explore if FOXO6 inhibits Wnt signaling in human cancers.

We additionally find that class I FOXs are candidate inhibitors of Wnt signaling. It was recently reported that conditional deletion of Foxi3 increased Wnt reporter activity in developing mouse teeth (*32*). Tooth formation requires tightly regulated Wnt pathway activity (*44*), suggesting that our identification of FOXIs as Wnt inhibitors is likely physiologically relevant. The mechanism by which FOXIs inhibit Wnt signaling requires further investigation. It has been suggested that FOXI1 decreases gastric cancer cell proliferation through the post-transcriptional destabilization of WNT3A mRNA (*45*). However, we find that at least in 293T cells, FOXI1 does not decrease WNT3A transcript levels, and that it inhibits Wnt signaling in the absence of Wnt co-receptors. Our results indicate that FOXIs may instead control β-catenin protein stability, as has been shown for FOXF2 (*46*).

The identification of FOXDs as candidate Wnt pathway activators is consistent with the recent observation that silencing of FOXD1 lowered β-catenin levels in prostate cancer cells, and concomitantly decreased cell proliferation and invasion (*31*). FOXD1-3 belong to a group of FOX proteins that strongly induce Wnt ligand expression, and we find that inhibition of WNT7/RECK signaling is sufficient to block the effect of FOXD1 in Wnt reporter assays. We have previously shown that FOXB2 controls Wnt activity in prostate cancer cells via WNT7/RECK (*28*), suggesting that these and potentially other FOX family members are functionally redundant in the context of Wnt signaling.

Although our study suggests substantial regulation of Wnt signaling by FOX transcription factors, it is important to note that these findings should be interpreted with due caution. It is known that the function of FOX proteins can differ considerably between different tissues and cell types. For example, it was recently shown that FOXF2 functions as a Wnt pathway inhibitor in basal-like breast cancer cells, but that it activates Wnt signaling in luminal breast cancer (*47*). The authors attributed these effects to the recruitment of specific transcription co-factors that differentially regulate the expression of Wnt ligands and receptors. Similarly, FOXQ1 has opposing functions in carcinoma and melanoma cells, which may be mediated by the cell type-dependent recruitment of β-catenin (*48*). Finally, it is likely that many FOX family members have more than one mode of action in the Wnt pathway as have been shown for, e.g., FOXM1 (*49*), FOXQ1 (*20*), and also FOXF2 (*46, 47, 50*). Thus, the relevance of our findings for different biological contexts necessitates thorough further investigation.

In conclusion, we provide a comprehensive overview of the function of FOX transcription factors in the Wnt signaling pathway. Similar to other transcription factor families, FOX proteins may act as context and gene-specific rheostats of Wnt pathway activity, and thereby fine-tune the transcriptional outcome of Wnt signaling. It is likely that these findings are important for cancer biology in particular.

## MATERIALS AND METHODS

### Cell lines

Authenticated 293T, HCT116, L, and L/Wnt3a cells were obtained from the German Collection of Microorganisms and Cell Cultures (DSMZ, Braunschweig, Germany) and the American Type Culture Collection (ATCC, Manassas, USA). 293T ΔCTNNB1, Penta-knockout and ΔLRP5/6 cells have been described elsewhere (*20, 21*). Cells were cultured in DMEM with 10% fetal bovine serum (FBS), 2 mM glutamine, and 1% penicillin and streptomycin at 37°C / 5% CO2. MCF-7 cells were kindly provided by Francisca Lottersberger (Linköping University, Sweden). All experiments were performed using low passage cells from confirmed mycoplasma-free frozen stocks. Wnt3a and control conditioned media were collected from stably transfected L cells, following the supplier’s guidelines. R-spondin 3 conditioned media were generated by transient transfection of Rspo3ΔC (*51*) into 293T cells.

### Plasmid construction and transfection

The construction and validation of the FOX transcription factor library used in this study has been described previously (*19*). The Flag-LEF1 plasmid and WNT7B promoter reporter were described in (*28*). For the WNT1 reporter, the WNT7B promoter was replaced by a 1.5kb fragment of the WNT1 promoter region. TurboID constructs were prepared by cloning of gene of interest into the N-terminally located TurboID pCS2 flag. All constructs were confirmed by partial sequencing (Eurofins Genomics, Ebersberg, Germany). Cell transfection was performed using Lipofectamine 2000 or Lipofectamine stem transfection reagent (Thermo Fisher Scientific, Waltham, USA), or jetOPTIMUS transfection reagents (Polyplus Transfection, Illkirch, France), according to the supplier’s recommendations. Other plasmids used in this study include: M50 Super 8x TOPflash and M51 Super 8x FOPflash ((*18*); Addgene plasmids 12456/12457, deposited by Randall Moon, University of Washington, School of Medicine, Seattle, USA); Renilla luciferase control plasmid (Addgene plasmid 12179, deposited by David Bartel, Whitehead Institute for Biomedical Research, Cambridge, USA); Flag-LRP6 ΔE1-4 ((*52*); a gift from Christof Niehrs, Institute of Molecular Biology, Mainz, Germany); and mCherry-Beta-Catenin-20 (Addgene plasmid 55001, deposited by Michael Davidson, Florida State University, Florida, USA). In some experiments, cells were additionally treated with CHIR90221 (Sigma Aldrich), LGK974 (Cayman Chemicals), or bortezomib (a gift from Padraig D’arcy, Linköping University). siRNAs against RECK (#1) and WNT7B (#3) were purchased from Integrated DNA Technologies (IDT, Skokie, USA), and validated previously (*28*). Scrambled siRNA control was from Thermo Fisher Scientific.

### Reporter assays

For reporter assays, cells were seeded on a 96-well plate and transfected with 50 ng firefly reporter, 5 ng renilla control, and 10 ng of the plasmid of interest in each well. Where indicated, 6 hours after transfection cells were treated with control, Wnt3a (standard dilution 1:5) or R-spondin 3 conditioned media (standard dilution 1:1,000). The dual luciferase assay was conducted as described previously (*53*) with few changes. Briefly, after overnight incubation, cells were lysed in passive lysis buffer (25 mM Tris, 2 mM DTT, 2 mM EDTA, 10% (v/v) glycerol, 1% (v/v) Triton X-100, (pH 7.8)) and agitated for 10 min. Lysates were transferred to a flat bottomed 96 well luminescence assay plate. Firefly luciferase buffer (200 μM D-luciferin in 200 mM Tris-HCl, 15 mM MgSO4, 100 μM EDTA, 1 mM ATP, 25 mM DTT, pH 8.0) was added to each well and the plate was incubated for 2 min at room temperature. Luciferase activity was measured using Spark10 (Tecan) or a SpectraMax iD3 Multi-Mode Microplate Reader (Molecular Devices). Next, Renilla luciferase buffer (4 μM coelenterazine-h in 500 mM NaCl, 500 mM Na2SO4, 10 mM NaOAc, 15 mM EDTA, 25 mM sodium pyrophosphate, 50 μM phenyl-benzothiazole, pH 5.0) was added to the plate and luminescence was measured immediately. Data were normalized to the Renilla control values, performed in triplicate.

### TaqMan qPCR array

qPCR array experiments were performed using custom 384-well TaqMan Gene Expression Array Cards (Thermo Fisher Scientific) containing inventoried probes for 48 gene targets, including three controls (18s rRNA, GAPDH, HPRT1). Data were normalized to the geometric mean of the controls. Experiments were done in 293T cells pretreated with 5 ng/ml recombinant human R-spondin 3 (R&D Systems, Minneapolis, USA). Experiments were performed with biological triplicates. For control of R-spondin effects, we included two untreated, empty transfected samples. Data were acquired on a QuantStudio 7 Flex Real-Time PCR system (Thermo Fisher Scientific). All gene expression data, excluding the untreated controls, can be found in supplemental Table 2.

### Quantitative realtime-PCR

qPCR was performed using standard protocols. Briefly, RNA was extracted using a Qiagen RNeasy mini kit (Hilden, Germany), and reverse transcribed using a high-capacity cDNA reverse transcription kit (Thermo Fisher Scientific). cDNA was amplified using validated custom primers, with SYBR green dye. Data were acquired on Bio-Rad CFX96 touch thermocycler (Hercules, USA) and normalized to HPRT1 control.

### Immunoblotting and immunoprecipitation

Cells were harvested in PBS and lysed in lysis buffer (1% NP-40 in PBS with 1 x protease inhibitor cocktail). Lysates were boiled in Laemmli sample buffer with 50 mM DTT, separated on 10% polyacrylamide gels (Bio-Rad, Hercules, USA), transferred onto nitrocellulose membranes, and incubated in blocking buffer (LI-COR, Lincoln, USA). Primary antibodies were detected using near-infrared (NIR) fluorophore-labelled secondary antibodies (LI-COR, Lincoln, USA). Blots were scanned on a LI-COR CLx imager. Antibodies used in this study are as follows: mouse anti-Flag M2 (F3165) and anti-Flag affinity gel (A2220) from Sigma Aldrich (St Louis, USA); rabbit anti-HSP70 (AF1663) from R&D Systems; rabbit anti-non phospho (Active) β-catenin (8814) from Cell Signaling Technology (Danvers, USA); mouse anti-FOXI (TA800145) from Thermo Fisher Scientific.

### TurboID sample preparation

The labelling and sample preparation of TurboID experiments was performed as described previously (*20, 22*). Briefly, N-terminal TurboID-FOX proteins and TurboID plasmids were transiently transfected into 293T cells using jetOPTIMUS (Polyplus Transfection). After 21 h of transfection, cells were treated with 50 μM biotin, and incubated for 3 h at 37 °C, 5% CO2. Cells were surface washed with ice cold PBS for three times to remove excess biotin and then harvested centrifuging at 1500 rpm for 15 min. Cells were washed thrice with ice-cold PBS buffer by centrifugation to remove any remaining biotin. Cells were lysed in RIPA buffer containing 1x protease inhibitor cocktail for 15 min on ice. Pre-washed streptavidin beads (GE Healthcare, USA) were added to the cell lysate and incubated overnight at 4 °C with end-over-end rotation. The beads were washed once with 1 ml of RIPA buffer, once with 1 ml of 1M KCl, once with 1ml of 0.1 M Na2CO3, once with 1 ml of 2 M urea in 10 mM Tris-HCl (pH 8.0), and twice with 1 ml RIPA lysis buffer. The beads then transferred to new Eppendorf tube and washed twice with 50 mM Tris-HCl buffer (pH 7.5) and 2 M urea/50 mM Tris (pH 7.5) buffer. Beads were incubated with 0.4 μg of trypsin (Thermo Fisher) in 2 M urea/50 mM Tris containing 1 mM DTT for 1 hour at 25 °C with end-over-end rotation. After incubation, the supernatant was collected and the beads were washed twice with 60 μl of 2 M urea/50 mM Tris buffer (pH 7.5) and the washes were combined with the collected supernatant. The supernatant was reduced with 4 mM DTT for 30 min at 25 °C with end-over-end rotation. The samples were alkylated with 10 mM iodoacetamide for 45 min in the dark at 25 °C with end-over-end rotation. For the complete digestion of the sample an additional 0.5 μg of trypsin was added and incubated at 25 °C overnight with end-over-end rotation. After overnight digestion the samples were desalted with C18 Thermo Fisher Scientific pipette tips and then dried with vacuum centrifuge.

### Mass spectrometry data acquisition and analysis

TurboID samples were analyzed by mass spectrometry, using an Easy nano LC 1200 system interfaced with a nanoEasy spray ion source (Thermo Fisher Scientific) connected Q Exactive HF Hybrid Quadrupole-Orbitrap Mass Spectrometer (Thermo Fisher Scientific). The peptides were loaded on a pre-column (Acclaim PepMap 100, 75 μm x 2 cm, Thermo FisherScientific) and the chromatographic separation was performed using an EASY-Spray C18 reversed-phase nano LC column (PepMap RSLC C18, 2 μm, 100A 75 μm x 25 cm, Thermo Fisher Scientific). The nanoLC was operating at 300 nl/min flow rate with a gradient (6-40% in 95 min and 5 min hold at 100%) solvent B (0.1% (v/v) formic acid in 100% acetonitrile) in solvent A (0.1% (v/v) formic acid in water) for 100 min. Separated peptides were electrosprayed and analyzed using a Q-Exactive HF mass spectrometer (Thermo Fisher Scientific), operated in positive polarity in a data-dependent mode. Full scans were performed at 120,000 resolutions at a range of 380–1400 m/z. The top 15 most intense multiple charged ions were isolated (1.2 m/z isolation window) and fragmented at a resolution of 30,000 with a dynamic exclusion of 30.0 s. Raw data were processed by Proteome Discover 2.0 (Thermo Fisher Scientific) searching against the *Homo sapiens* UniProt database (release 2019-12-16) with Sequest HT search engine. The search parameters were: Taxonomy: *Homo sapiens;* Enzymes; trypsin with two missed cleavages, no variable Modifications; fixed modification: Carbamidomethyl; Peptide Mass Tolerance, 10 ppm; MS/MS Fragment Tolerance, 0.02 Da. Quantification of the analyzed data were performed with Scaffold 5.1.0 (Proteome Software Inc., Portland, OR, USA) using total spectral count. Protein and peptide identifications were accepted if they could be established at greater than 95 and 90% probability, respectively and if protein identification contained at least two identified peptides.

### Colony formation assays

HCT116 were seeded on 6 well plates at a density of 1,000 cells after 24 hours of transfection. Cells were grown for 14 days. At the end of the experiment, cells were washed with PBS, and stained with 2% methylene blue in 50% methanol. Stained plates were photographed and colonies were counted manually. Colonies with less than 50 cells were excluded.

### Data quantification and analysis

Most statistical analyses were performed in R v4.2.0 (*54*), and were based on three or more biological replicates per condition. Statistical tests are indicated in the figure legends. Hierarchical clustering was done using the kmeans and hclust functions implemented in R package ComplexHeatmaps v2.12.1 (*55*), with a fixed seed and 1,000 iterations. Apart from data in figure 1, optimal cluster numbers were determined using the Silhouette method (*56*) implemented in R package factoextra v1.0.7 (*57*). Proteomics data were processed using SAINTexpress, CRAPome, and ProHits-viz (*58–60*), as described previously (*20*), including data from (*27*) following mouse-to-human gene name conversion. Gene ontology and enrichment analyses of proteomics data were performed using R package clusterProfiler v4.4.4 (*61*). CORUM protein complexes v4.1 have been described in (*26*). Proteomics data in supplemental Fig. 4 are from (*62*), their supplemental Table S3. Only hits with a spectral count >1 were included in the analysis. Bar graphs depict group means with standard deviation and individual data points.

## Data availability

TurboID proteomics data have been deposited at the PRIDE repository, and will be made available at the time of publication. Taqman qPCR array data can be found in supplemental Table 2.

## Acknowledgements

The authors thank Giulia Pizzolato, Francisca Lottersberger, Padraig D’arcy, Claudio Cantù and their teams for cells, reagents, and comments on the project. S. K. is a Wallenberg Molecular Medicine fellow, and acknowledges long-term support from the Knut and Alice Wallenberg Foundation. Additional funding for research in the Koch laboratory came from the Swedish Research Council (VR) and the Swedish Cancer Society (Cancerfonden). We thank the staff at the Core Facilities of the Faculty of Medicine and Health Sciences at Linköping University for mass spectrometry and molecular biology support.

---

**Supplemental figure 1:**
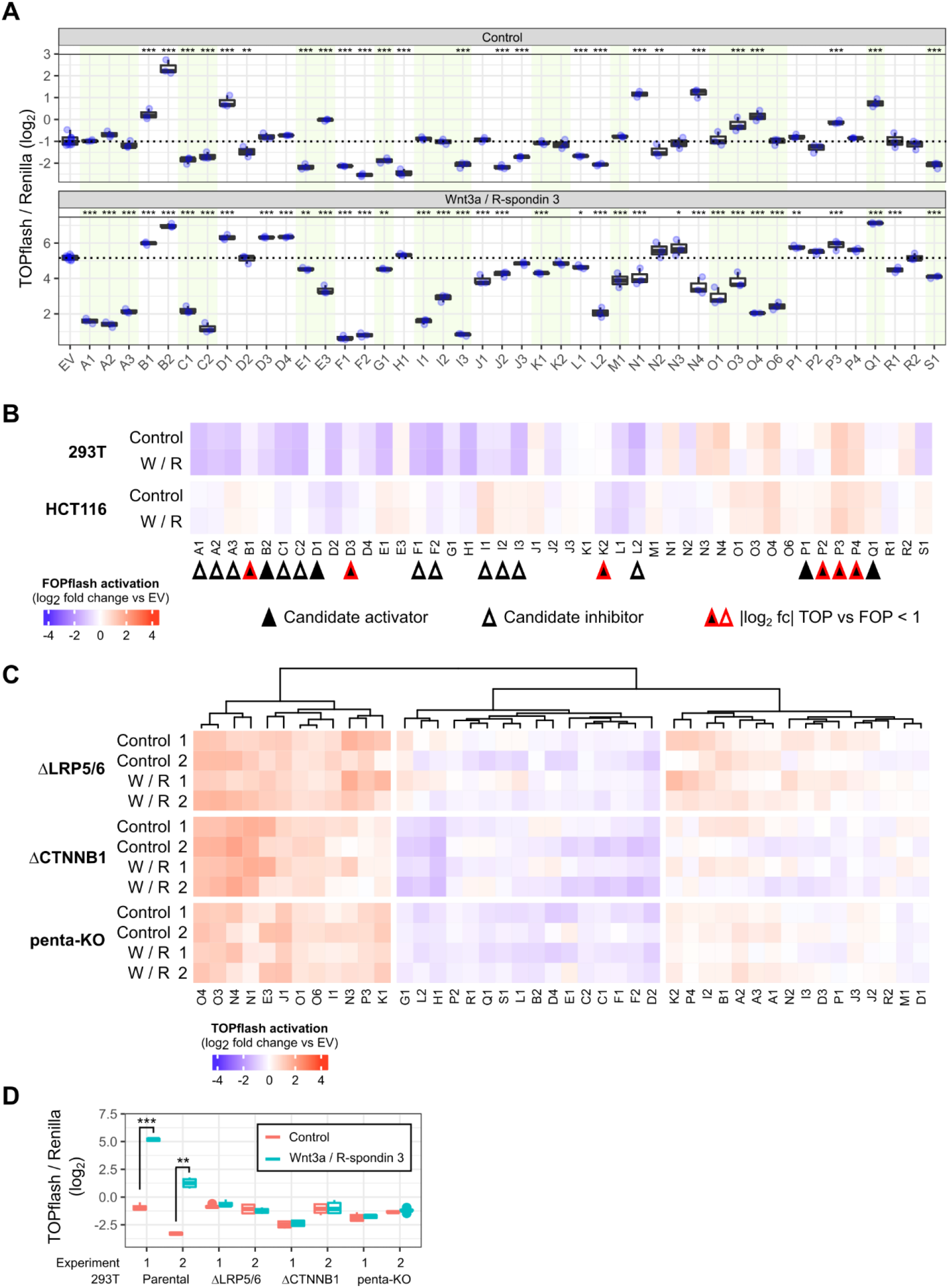
TOPflash assay controls. (A) Representative TOPflash results with individual data points, corresponding to 293T experiment 1 in Fig. 1B. Data are displayed as the Firefly to Renilla luciferase activity without additional normalization to empty vector (EV) control. Statistical significance was determined by Dunnett’s post-hoc test following one-way ANOVA. (B) Activity assay using the control reporter plasmid FOPflash, which contains scrambled TCF binding sites. Data were normalized to EV control. Where indicated, cells were treated with Wnt3a / R-spondin 3 (W/R) conditioned media. Candidate activators and inhibitors were defined in Fig. 1D. For candidates marked with a red outline, we observed a smaller than 2-fold difference in TOPflash versus FOPflash regulation. (C) TOPflash assay in 293T lacking the Wnt co-receptors LRP5/6 (ΔLRP5/6), β-catenin (ΔCTNNB1), or β-catenin and all TCF/LEF proteins (penta-KO). (D) TOPflash results from all EV-transfected cells used in the previous assays, following Wnt3a / R-spondin 3 treatment. Statistical significance in panel A was determined by Dunnett’s post-hoc test following one-way ANOVA. Data in B were analyzed using an unpaired Welch’s t-test with Bonferroni-Hochberg correction for multiple testing. *** *P*<0.001, ** *P*<0.01, * *P*<0.05.

**Supplemental figure 2:**
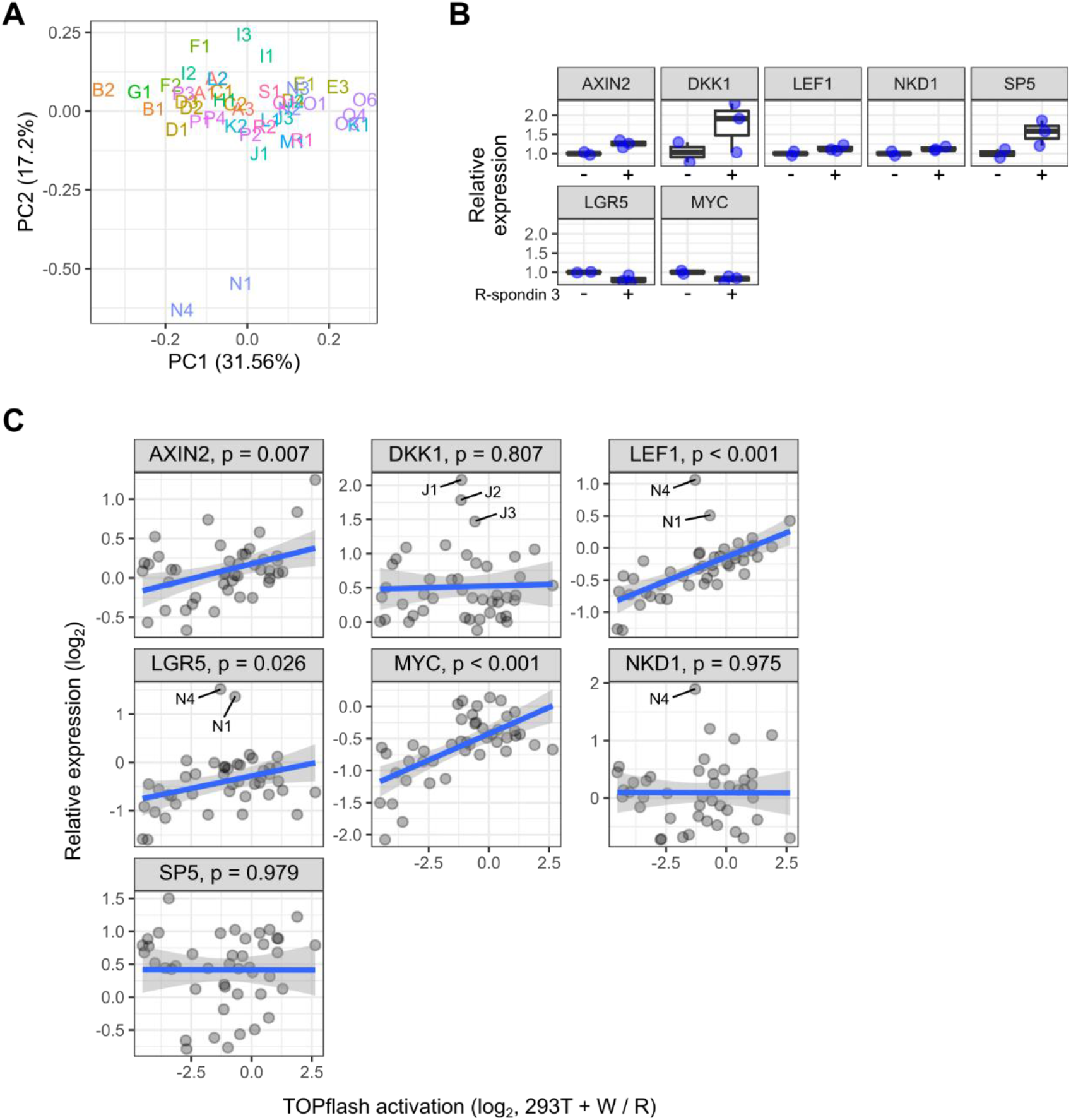
qPCR array controls. (A) Principal component plot based on expression changes of all genes included in the qPCR array, following normalization to housekeeping controls. (B) Expression changes of Wnt target genes included in the array following treatment with 5 ng/ml recombinant human R-spondin 3 (n=3) versus untreated control (n=2). All cells were transfected with empty vector. (C) Correlation analyses of Wnt target gene expression changes against averaged results from TOPflash assays in Fig. 1B. Notable outliers are highlighted. *P*-values are based on Spearman’s rank correlation.

**Supplemental figure 3:**
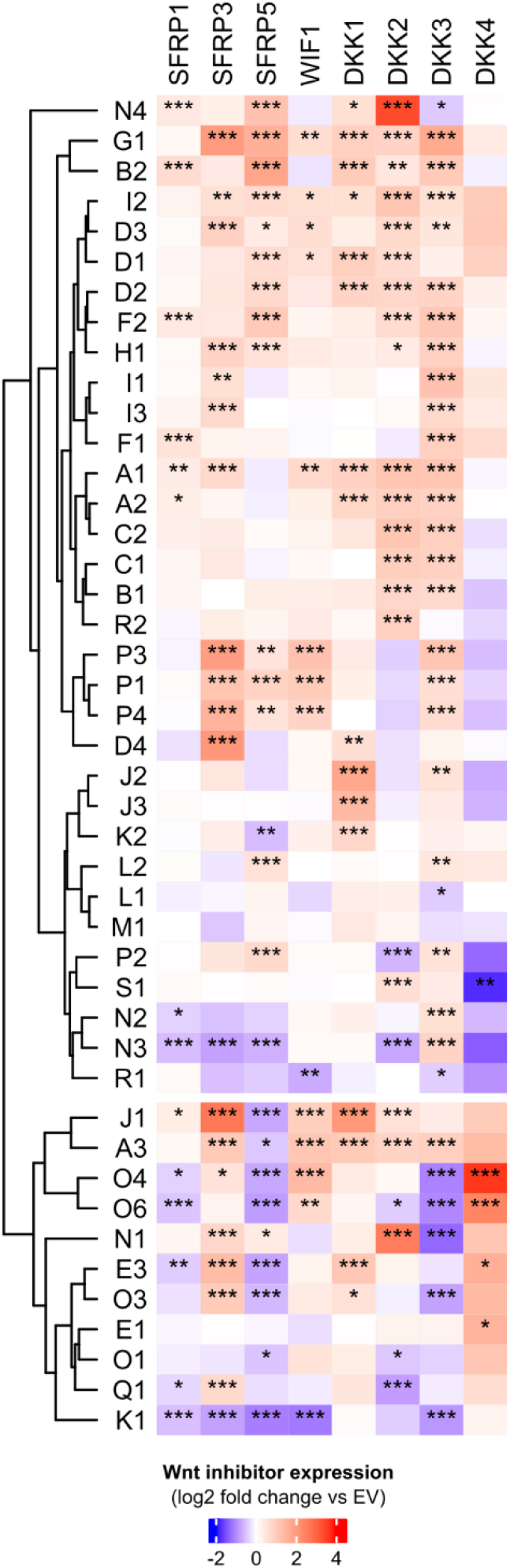
FOX proteins regulate secreted Wnt pathway inhibitors. (A) qPCR array of secreted Wnt pathway inhibitors in 293T treated with 5 ng/ml recombinant human R-spondin 3. Each cell represents the average of three biological replicates, normalized to empty vector (EV) control. Note that SFRP2 was beyond the limit of detection in most samples, and was therefore omitted from this analysis. Data for DKK1 are repeated from Wnt target gene analyses in Fig. 1C. Data were analyzed using Dunnett’s post-hoc test against EV following one-way ANOVA (*** *P*<0.001, ** *P*<0.01, * *P*<0.05).

**Supplemental figure 4:**
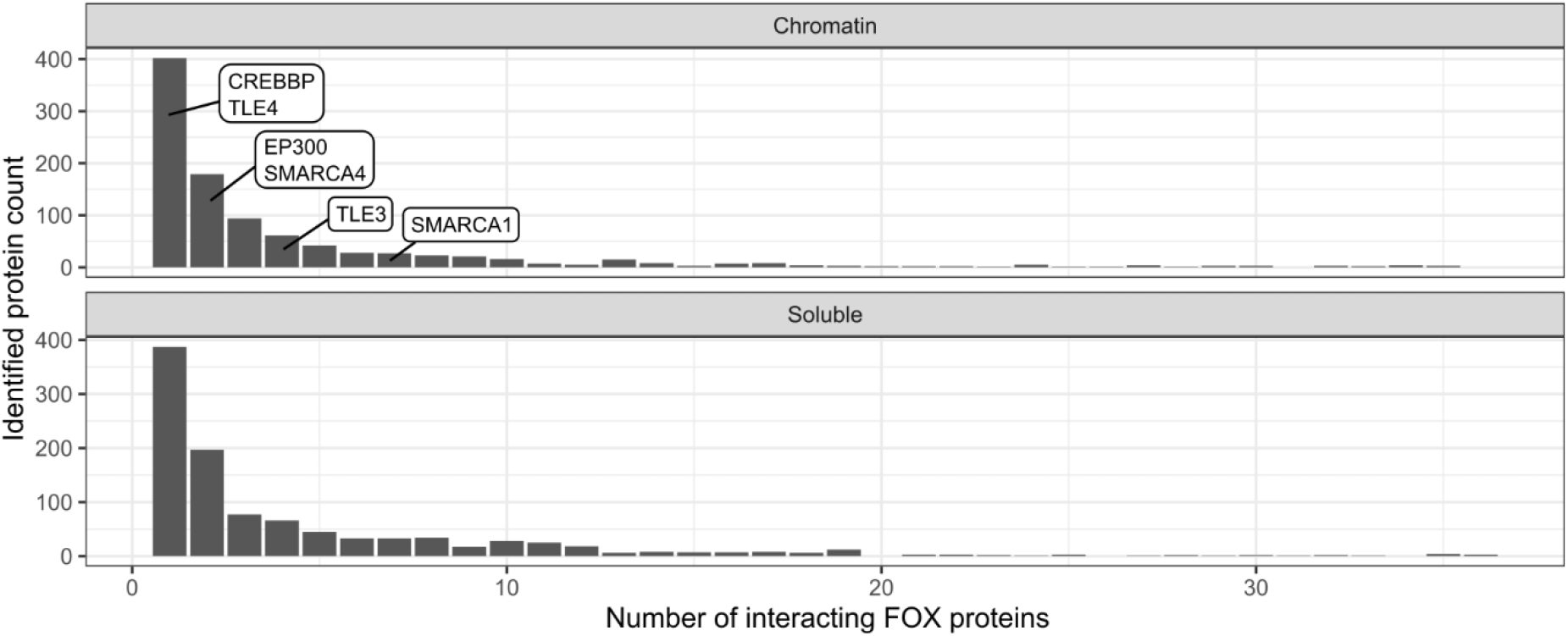
Overview of potential shared FOX interactors. Candidate FOX interactors identified by Li et al. (*24*) using co-immunoprecipitation / mass spectrometry following expression of 36 FOX family members in 293T. Data are separated into chromatin and soluble sample fractions. Some interactors of interest that were also found in our TurboID data are highlighted.

**Supplemental figure 5:**
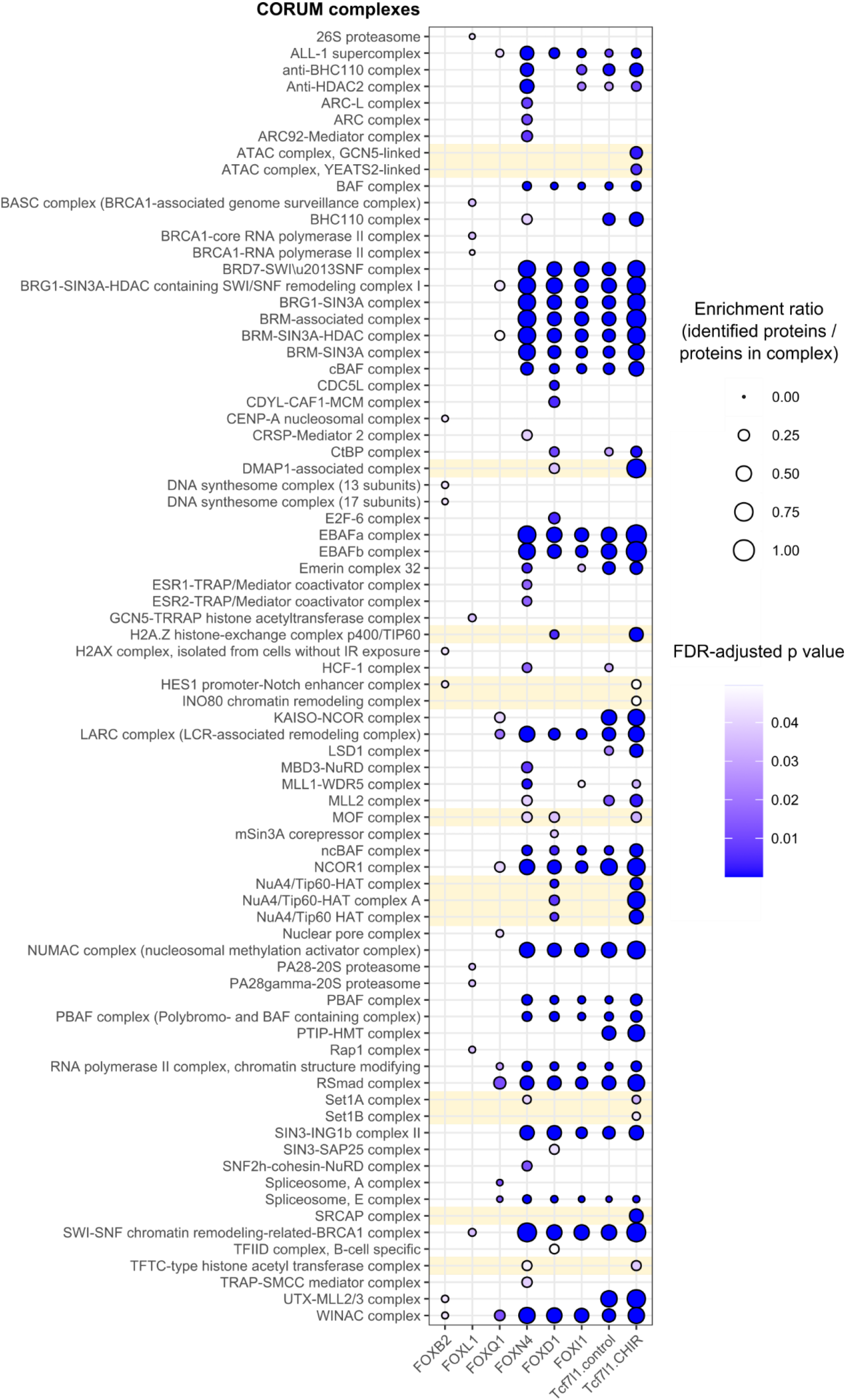
FOX proteins and Tcf7l1 share interacting protein complexes. Enrichment analysis against all human protein complexes curated in the CORUM database (*26*). All complexes that were significantly enriched in at least one FOX sample or Tcf7l1 are shown. Results include proteomics data from earlier BioID-FOXB2, TurboID-FOXQ1, and BioID-Tcf7l1 studies (*20, 27, 28*). Note that no enriched complexes were identified for FOXG1. Complex names were taken from the CORUM resource as-is, and contain broadly overlapping complexes. Yellow highlighting indicates protein complexes that are associated with Tcf7l1 after CHIR treatment. FDR, false discovery rate.

**Supplemental figure 6:**
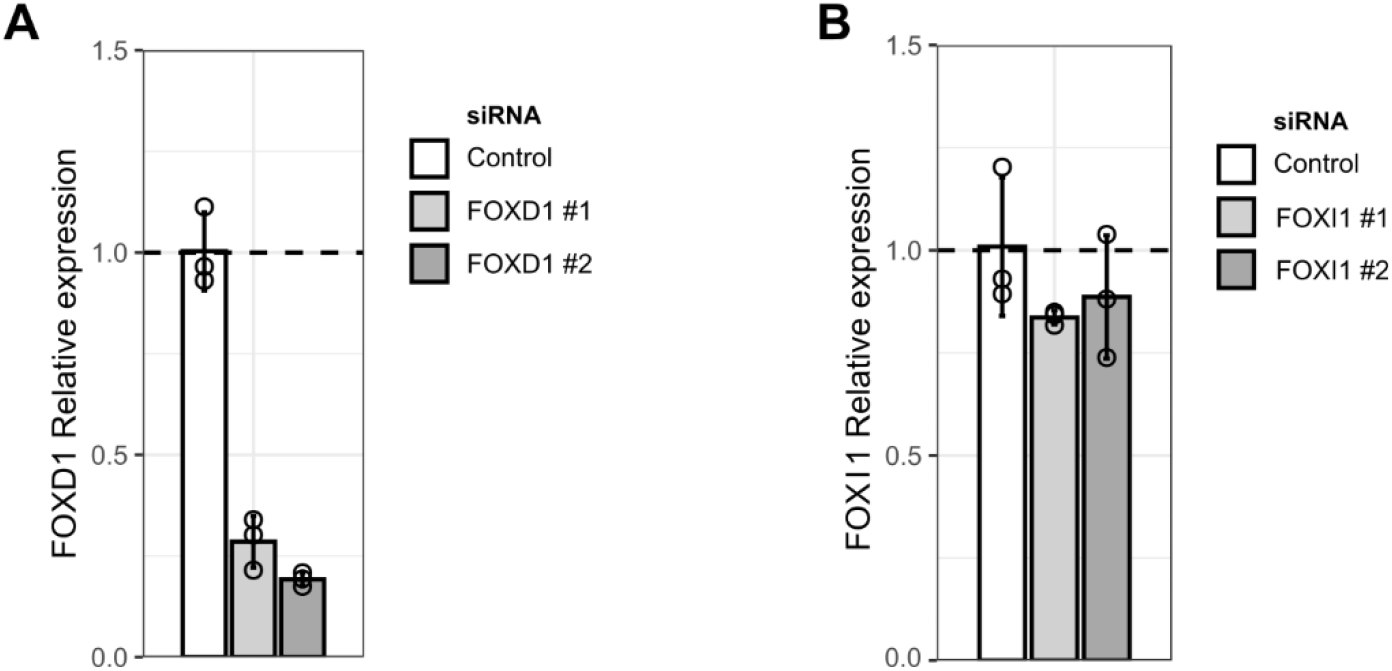
Control of FOXD1 and FOXI1 knock-down efficiency. (A, B) qPCR analysis of (A) FOXD1 and (B) FOXI expression in 293T, following RNA interference using two independent siRNAs.

**Supplemental figure 7:**
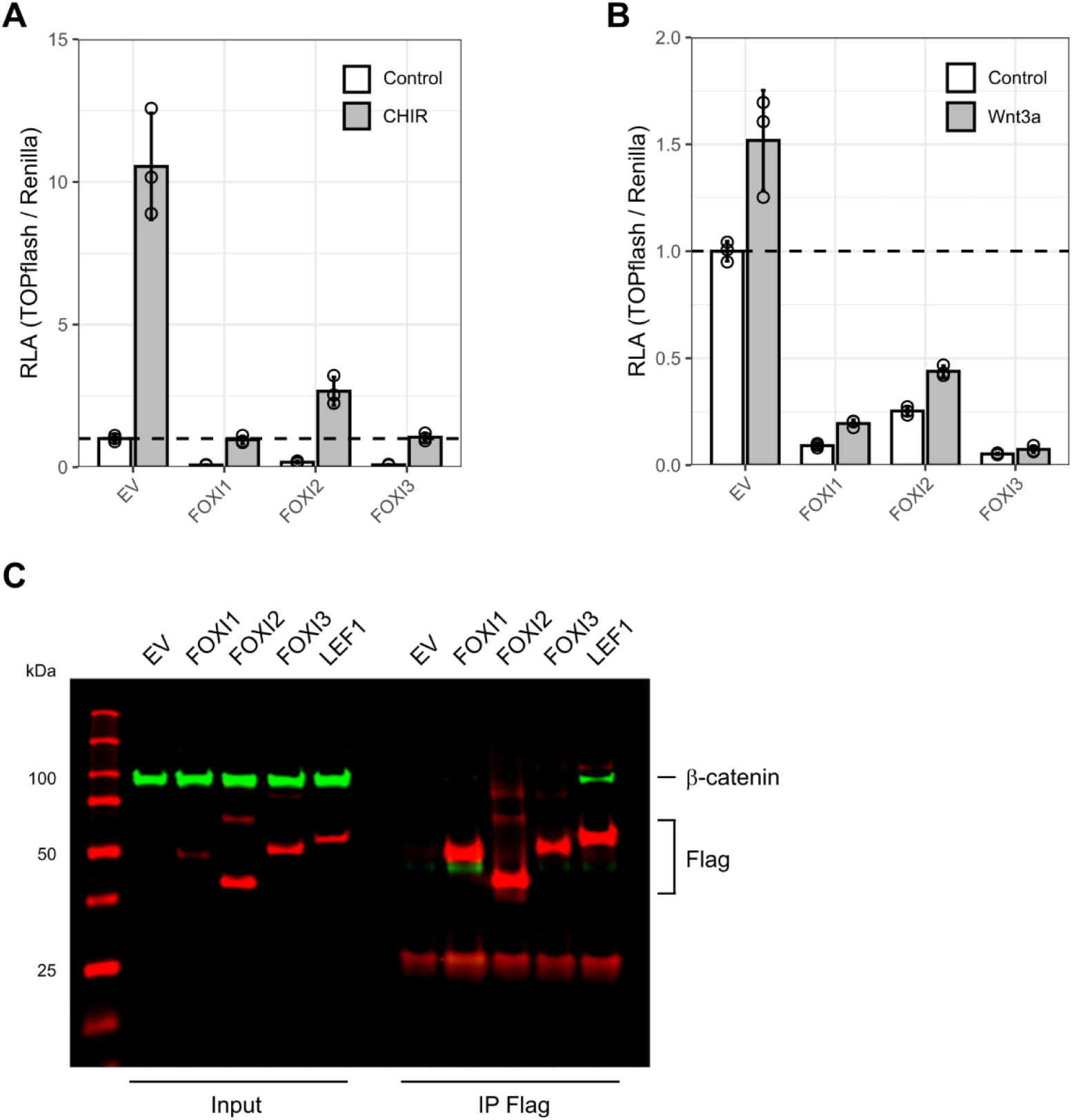
FOXIs inhibit β-catenin signaling. (A) TOPflash assay in HCT116 following treatment with 5 μM GSK3 inhibitor CHIR99021. (B) TOPflash assay in 293T following β-catenin overexpression. Where indicated, cells were treated with Wnt3a conditioned media. (C) Co-immunoprecipitation assay in HCT116. The indicated proteins were precipitated using anti-Flag agarose. LEF1 was included as a positive control. RLA, relative luciferase activity.

**Supplemental figure 8:**
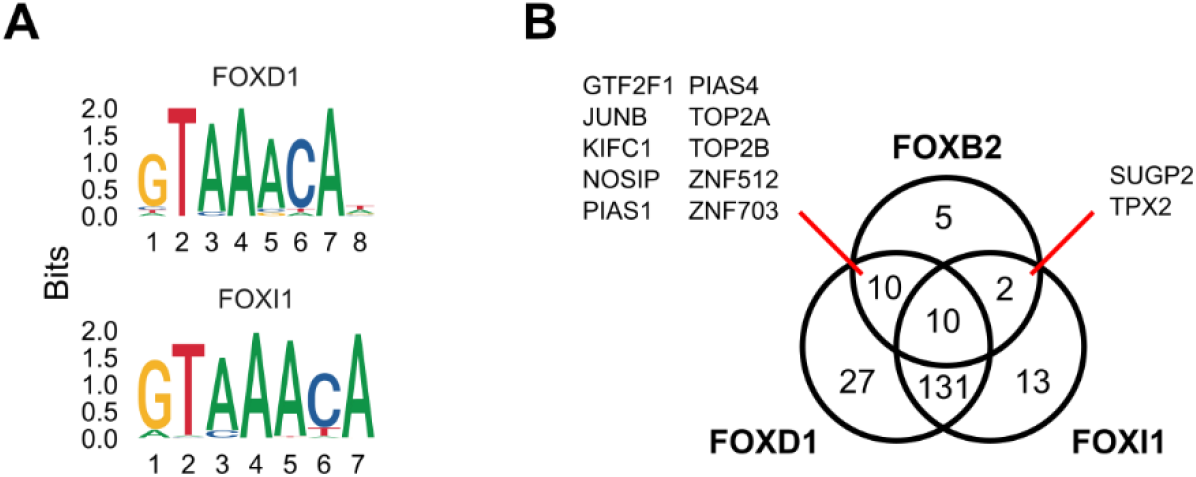
FOXD1 and FOXI1 have distinct interactors. (A) Position-weighted matrices of FOXD1 (MA0031.1) and FOXI1 (MA0042.2), as curated in the JASPAR 2022 database (*63*). (B) Shared interactors between FOXD1, FOXI1, and FOXB2, based on proximity proteomics.

## Notes

### Competing Interest Statement

The authors have declared no competing interest.

## REFERENCES

1. R. Nusse, H. Clevers, Wnt/β-catenin signaling, disease, and emerging therapeutic modalities. Cell 169, 985–999 (2017).

2. A. Franz, D. Shlyueva, E. Brunner, A. Stark, K. Basler, Probing the canonicity of the Wnt/Wingless signaling pathway. PLOS Genetics 13, e1006700 (2017).

3. J. Schuijers, M. Mokry, P. Hatzis, E. Cuppen, H. Clevers, Wnt-induced transcriptional activation is exclusively mediated by TCF/LEF. The EMBO journal 33, 146–156 (2014).

4. D. Zimmerli, C. Borrelli, A. Jauregi-Miguel, S. Soderholm, S. Brutsch, N. Doumpas, J. Reichmuth, F. Murphy-Seiler, M. Aguet, K. Basler, A. E. Moor, C. Cantu, TBX3 acts as tissue-specific component of the Wnt/beta-catenin transcriptional complex. Elife 9, e58123 (2020).

5. A.-B. Ramakrishnan, L. Chen, Peter E. Burby, Ken M. Cadigan, Wnt target enhancer regulation by a CDX/TCF transcription factor collective and a novel DNA motif. Nucleic Acids Research 49, 8625–8641 (2021).

6. A. B. Ramakrishnan, P. E. Burby, K. Adiga, K. M. Cadigan, SOX9 and TCF transcription factors associate to mediate Wnt/beta-catenin target gene activation in colorectal cancer. J Biol Chem, 102735 (2022).

7. M. L. Golson, K. H. Kaestner, Fox transcription factors: from development to disease. Development 143, 4558–4570 (2016).

8. E. W. F. Lam, J. J. Brosens, A. R. Gomes, C.-Y. Koo, Forkhead box proteins: tuning forks for transcriptional harmony. Nature Reviews Cancer 13, 482–495 (2013).

9. L. Herman, A.-L. Todeschini, R. A. Veitia, Forkhead Transcription Factors in Health and Disease. Trends in Genetics 37, 460–475 (2021).

10. J.-H. Paik, R. Kollipara, G. Chu, H. Ji, Y. Xiao, Z. Ding, L. Miao, Z. Tothova, J. W. Horner, D. R. Carrasco, S. Jiang, D. G. Gilliland, L. Chin, W. H. Wong, D. H. Castrillon, R. A. DePinho, FoxOs Are Lineage-Restricted Redundant Tumor Suppressors and Regulate Endothelial Cell Homeostasis. Cell 128, 309–323 (2007).

11. S. M. Ahmad, P. Bhattacharyya, N. Jeffries, S. S. Gisselbrecht, A. M. Michelson, Two Forkhead transcription factors regulate cardiac progenitor specification by controlling the expression of receptors of the fibroblast growth factor and Wnt signaling pathways. Development 143, 306–317 (2016).

12. L. Molin, A. Mounsey, S. Aslam, P. Bauer, J. Young, M. James, A. Sharma-Oates, I. A. Hope, Evolutionary conservation of redundancy between a diverged pair of forkhead transcription factor homologues. Development 127, 4825–4835 (2000).

13. M. Ormestad, J. Astorga, H. Landgren, T. Wang, B. R. Johansson, N. Miura, P. Carlsson, Foxf1 and Foxf2 control murine gut development by limiting mesenchymal Wnt signaling and promoting extracellular matrix production. Development 133, 833–843 (2006).

14. B. A. Benayoun, S. Caburet, R. A. Veitia, Forkhead transcription factors: key players in health and disease. Trends in Genetics 27, 224–232 (2011).

15. S. Koch, Regulation of Wnt Signaling by FOX Transcription Factors in Cancer. Cancers 13, 3446 (2021).

16. A. Gong, S. Huang, FoxM1 and Wnt/β-Catenin Signaling in Glioma Stem Cells. Cancer Research 72, 5658–5662 (2012).

17. M. A. Essers, L. M. de Vries-Smits, N. Barker, P. E. Polderman, B. M. Burgering, H. C. Korswagen, Functional interaction between β-catenin and FOXO in oxidative stress signaling. Science 308, 1181–1184 (2005).

18. M. T. Veeman, D. C. Slusarski, A. Kaykas, S. H. Louie, R. T. Moon, Zebrafish prickle, a modulator of noncanonical Wnt/Fz signaling, regulates gastrulation movements. Current biology 13, 680–685 (2003).

19. L. Moparthi, S. Koch, A uniform expression library for the exploration of FOX transcription factor biology. Differentiation 115, 30–36 (2020).

20. G. Pizzolato, L. Moparthi, S. Soderholm, C. Cantu, S. Koch, The oncogenic transcription factor FOXQ1 is a differential regulator of Wnt target genes. J Cell Sci 135, jcs.260082 (2022).

21. N. Doumpas, F. Lampart, M. D. Robinson, A. Lentini, C. E. Nestor, C. Cantu, K. Basler, TCF/LEF dependent and independent transcriptional regulation of Wnt/beta-catenin target genes. EMBO J 38, (2019).

22. T. C. Branon, J. A. Bosch, A. D. Sanchez, N. D. Udeshi, T. Svinkina, S. A. Carr, J. L. Feldman, N. Perrimon, A. Y. Ting, Efficient proximity labeling in living cells and organisms with TurboID. Nature biotechnology 36, 880–887 (2018).

23. N. Alerasool, H. Leng, Z.-Y. Lin, A.-C. Gingras, M. Taipale, Identification and functional characterization of transcriptional activators in human cells. Molecular Cell 82, 677–695.e677 (2022).

24. X. Li, W. Wang, J. Wang, A. Malovannaya, Y. Xi, W. Li, R. Guerra, D. H. Hawke, J. Qin, J. Chen, Proteomic analyses reveal distinct chromatin-associated and soluble transcription factor complexes. Mol Syst Biol 11, 775 (2015).

25. J.-P. Lambert, M. Tucholska, C. Go, J. D. Knight, A.-C. Gingras, Proximity biotinylation and affinity purification are complementary approaches for the interactome mapping of chromatin-associated protein complexes. Journal of proteomics 118, 81–94 (2015).

26. G. Tsitsiridis, R. Steinkamp, M. Giurgiu, B. Brauner, G. Fobo, G. Frishman, C. Montrone, A. Ruepp, CORUM: the comprehensive resource of mammalian protein complexes–2022. Nucleic Acids Research, (2022).

27. S. Moreira, C. Seo, V. Gordon, S. Xing, R. Wu, E. Polena, V. Fung, D. Ng, C. J. Wong, B. Larsen, B. Raught, A.-C. Gingras, Y. Lu, B. W. Doble, Endogenous BioID elucidates TCF7L1 interactome modulation upon GSK-3 inhibition in mouse ESCs. bioRxiv, 431023 (2018).

28. L. Moparthi, G. Pizzolato, S. Koch, Wnt activator FOXB2 drives the neuroendocrine differentiation of prostate cancer. Proc Natl Acad Sci U S A 116, 22189–22195 (2019).

29. S. Söderholm, C. Cantù, The WNT/β-catenin dependent transcription: A tissue-specific business. WIREs Mechanisms of Disease 13, e1511 (2021).

30. C. Wan, S. Mahara, C. Sun, A. Doan, H. K. Chua, D. Xu, J. Bian, Y. Li, D. Zhu, D. Sooraj, T. Cierpicki, J. Grembecka, R. Firestein, Genome-scale CRISPR-Cas9 screen of Wnt/beta-catenin signaling identifies therapeutic targets for colorectal cancer. Sci Adv 7, eabf2567 (2021).

31. C. Donmez, E. Konac, Silencing effects of FOXD1 inhibit metastatic potentials of the PCa via N-cadherin - Wnt/beta-catenin crosstalk. Gene 836, 146680 (2022).

32. I. Mogollón, N. Kangasniemi, J. E. Moustakas-Verho, L. Ahtiainen, Foxi3 Suppresses Signaling Center Fate and is Necessary for the Early Development of Mouse Teeth. bioRxiv, 2022.2007.2018.500404 (2022).

33. T. W. Li, J. H. Ting, N. N. Yokoyama, A. Bernstein, M. van de Wetering, M. L. Waterman, Wnt activation and alternative promoter repression of LEF1 in colon cancer. Mol Cell Biol 26, 5284–5299 (2006).

34. J. Fröhlich, K. Rose, A. Hecht, Transcriptional activity mediated by β-CATENIN and TCF/LEF family members is completely dispensable for survival of multiple human colorectal cancer cell lines. bioRxiv, 2022.2012.2005.519142 (2022).

35. D. Hoogeboom, M. A. Essers, P. E. Polderman, E. Voets, L. M. Smits, B. M. Burgering, Interaction of FOXO with beta-catenin inhibits beta-catenin/T cell factor activity. J Biol Chem 283, 9224–9230 (2008).

36. H. Yamamoto, M. Ihara, Y. Matsuura, A. Kikuchi, Sumoylation is involved in beta-catenin-dependent activation of Tcf-4. EMBO J 22, 2047–2059 (2003).

37. J. Xue, Y. Chen, Y. Wu, Z. Wang, A. Zhou, S. Zhang, K. Lin, K. Aldape, S. Majumder, Z. Lu, S. Huang, Tumour suppressor TRIM33 targets nuclear β-catenin degradation. Nature Communications 6, 6156 (2015).

38. X. Xia, F. Zuo, M. Luo, Y. Sun, J. Bai, Q. Xi, Role of TRIM33 in Wnt signaling during mesendoderm differentiation. Science China Life Sciences 60, 1142–1149 (2017).

39. A. Sundqvist, M. Morikawa, J. Ren, E. Vasilaki, N. Kawasaki, M. Kobayashi, D. Koinuma, H. Aburatani, K. Miyazono, C. H. Heldin, H. van Dam, P. Ten Dijke, JUNB governs a feed-forward network of TGFbeta signaling that aggravates breast cancer invasion. Nucleic Acids Res 46, 1180–1195 (2018).

40. J.-h. Paik, Z. Ding, R. Narurkar, S. Ramkissoon, F. Muller, W. S. Kamoun, S.-S. Chae, H. Zheng, H. Ying, J. Mahoney, D. Hiller, S. Jiang, A. Protopopov, W. H. Wong, L. Chin, K. L. Ligon, R. A. DePinho, FoxOs Cooperatively Regulate Diverse Pathways Governing Neural Stem Cell Homeostasis. Cell Stem Cell 5, 540–553 (2009).

41. F. M. Jacobs, L. P. van der Heide, P. J. Wijchers, J. P. Burbach, M. F. Hoekman, M. P. Smidt, FoxO6, a novel member of the FoxO class of transcription factors with distinct shuttling dynamics. J Biol Chem 278, 35959–35967 (2003).

42. Q. Li, H. Tang, F. Hu, C. Qin, Silencing of FOXO6 inhibits the proliferation, invasion, and glycolysis in colorectal cancer cells. J Cell Biochem 120, 3853–3860 (2019).

43. F. Lallemand, A. Petitalot, S. Vacher, L. de Koning, K. Taouis, B. S. Lopez, S. Zinn-Justin, N. Dalla-Venezia, W. Chemlali, A. Schnitzler, R. Lidereau, I. Bieche, S. M. Caputo, Involvement of the FOXO6 transcriptional factor in breast carcinogenesis. Oncotarget 9, 7464–7475 (2018).

44. E. Jarvinen, I. Salazar-Ciudad, W. Birchmeier, M. M. Taketo, J. Jernvall, I. Thesleff, Continuous tooth generation in mouse is induced by activated epithelial Wnt/beta-catenin signaling. Proc Natl Acad Sci U S A 103, 18627–18632 (2006).

45. R. Sun, Z. Liu, D. Tong, Y. Yang, B. Guo, X. Wang, L. Zhao, C. Huang, miR-491-5p, mediated by Foxi1, functions as a tumor suppressor by targeting Wnt3a/β-catenin signaling in the development of gastric cancer. Cell Death & Disease 8, e2714–e2714 (2017).

46. A. Higashimori, Y. Dong, Y. Zhang, W. Kang, G. Nakatsu, S. S. M. Ng, T. Arakawa, J. J. Y. Sung, F. K. L. Chan, J. Yu, Forkhead Box F2 Suppresses Gastric Cancer through a Novel FOXF2-IRF2BPL-beta-Catenin Signaling Axis. Cancer Res 78, 1643–1656 (2018).

47. X. Zhang, R. Zhang, C. Hou, R. He, Q. S. Wang, T. H. Zhou, X. Q. Li, Q. L. Zhai, Y. M. Feng, FOXF2 oppositely regulates stemness in luminal and basal-like breast cancer cells through the Wnt/beta-catenin pathway. J Biol Chem 298, 102082 (2022).

48. A. Bagati, A. Bianchi-Smiraglia, S. Moparthy, K. Kolesnikova, E. E. Fink, B. C. Lipchick, M. Kolesnikova, P. Jowdy, A. Polechetti, A. Mahpour, J. Ross, J. A. Wawrzyniak, D. H. Yun, G. Paragh, N. I. Kozlova, A. E. Berman, J. Wang, S. Liu, M. J. Nemeth, M. A. Nikiforov, Melanoma Suppressor Functions of the Carcinoma Oncogene FOXQ1. Cell Reports 20, 2820–2832 (2017).

49. N. Zhang, P. Wei, A. Gong, W. T. Chiu, H. T. Lee, H. Colman, H. Huang, J. Xue, M. Liu, Y. Wang, R. Sawaya, K. Xie, W. K. Yung, R. H. Medema, X. He, S. Huang, FoxM1 promotes beta-catenin nuclear localization and controls Wnt target-gene expression and glioma tumorigenesis. Cancer Cell 20, 427–442 (2011).

50. A. M. Nik, A. Reyahi, F. Ponten, P. Carlsson, Foxf2 in intestinal fibroblasts reduces numbers of Lgr5(+) stem cells and adenoma formation by inhibiting Wnt signaling. Gastroenterology 144, 1001–1011 (2013).

51. B. Ohkawara, A. Glinka, C. Niehrs, Rspo3 binds syndecan 4 and induces Wnt/PCP signaling via clathrin-mediated endocytosis to promote morphogenesis. Developmental cell 20, 303–314 (2011).

52. G. Davidson, W. Wu, J. Shen, J. Bilic, U. Fenger, P. Stannek, A. Glinka, C. Niehrs, Casein kinase 1 gamma couples Wnt receptor activation to cytoplasmic signal transduction. Nature 438, 867–872 (2005).

53. M. Hampf, M. Gossen, A protocol for combined Photinus and Renilla luciferase quantification compatible with protein assays. Analytical biochemistry 356, 94–99 (2006).

54. R Core Team. (R Foundation for Statistical Computing, Vienna, Austria, 2022).

55. Z. Gu, R. Eils, M. Schlesner, Complex heatmaps reveal patterns and correlations in multidimensional genomic data. Bioinformatics 32, 2847–2849 (2016).

56. P. J. Rousseeuw, Silhouettes: A graphical aid to the interpretation and validation of cluster analysis. Journal of Computational and Applied Mathematics 20, 53–65 (1987).

57. A. Kassambara, F. Mundt. (2020).

58. G. Teo, G. Liu, J. Zhang, A. I. Nesvizhskii, A.-C. Gingras, H. Choi, SAINTexpress: Improvements and additional features in Significance Analysis of INTeractome software. Journal of Proteomics 100, 37–43 (2014).

59. D. Mellacheruvu, Z. Wright, A. L. Couzens, J.-P. Lambert, N. A. St-Denis, T. Li, Y. V. Miteva, S. Hauri, M. E. Sardiu, T. Y. Low, V. A. Halim, R. D. Bagshaw, N. C. Hubner, A. al-Hakim, A. Bouchard, D. Faubert, D. Fermin, W. H. Dunham, M. Goudreault, Z.-Y. Lin, B. G. Badillo, T. Pawson, D. Durocher, B. Coulombe, R. Aebersold, G. Superti-Furga, J. Colinge, A. J. R. Heck, H. Choi, M. Gstaiger, S. Mohammed, I. M. Cristea, K. L. Bennett, M. P. Washburn, B. Raught, R. M. Ewing, A.-C. Gingras, A. I. Nesvizhskii, The CRAPome: a contaminant repository for affinity purification–mass spectrometry data. Nature Methods 10, 730–736 (2013).

60. J. D. R. Knight, H. Choi, G. D. Gupta, L. Pelletier, B. Raught, A. I. Nesvizhskii, A.-C. Gingras, ProHits-viz: a suite of web tools for visualizing interaction proteomics data. Nature Methods 14, 645–646 (2017).

61. T. Wu, E. Hu, S. Xu, M. Chen, P. Guo, Z. Dai, T. Feng, L. Zhou, W. Tang, L. Zhan, X. Fu, S. Liu, X. Bo, G. Yu, clusterProfiler 4.0: A universal enrichment tool for interpreting omics data. The Innovation 2, 100141 (2021).

62. X. Li, W. Wang, J. Wang, A. Malovannaya, Y. Xi, W. Li, R. Guerra, D. H. Hawke, J. Qin, J. Chen, Proteomic analyses reveal distinct chromatin-associated and soluble transcription factor complexes. Molecular Systems Biology 11, 775 (2015).

63. J. A. Castro-Mondragon, R. Riudavets-Puig, I. Rauluseviciute, R. Berhanu Lemma, L. Turchi, R. Blanc-Mathieu, J. Lucas, P. Boddie, A. Khan, N. Manosalva Pérez, O. Fornes, Tiffany Y. Leung, A. Aguirre, F. Hammal, D. Schmelter, D. Baranasic, B. Ballester, A. Sandelin, B. Lenhard, K. Vandepoele, W. W. Wasserman, F. Parcy, A. Mathelier, JASPAR 2022: the 9th release of the open-access database of transcription factor binding profiles. Nucleic Acids Research 50, D165–D173 (2021).

